# Moderate-intensity exercise might be a competitive protocol to prevent obesity and obesity-related metabolic diseases

**DOI:** 10.1101/2021.11.18.469147

**Authors:** Ryeo-Won Kwon, Seung-Jea Lee, Tae-Wook Kim, Jeong Beom Lee

## Abstract

To determine effects of exercise intensity on expression levels of cytokines and neurotransmitters beneficial for the prevention and improvement of obesity and obesity-related metabolic diseases. Expression levels of irisin, fibroblast growth factor-21 (FGF21), adiponectin, free fatty acid (FFA), dopamine (DA), and serotonin (5-HT) levels after moderate-intensity (50% of maximal oxygen uptake (VO_2_max)) and high-intensity treadmill running (80% VO_2_max) for 30 min in 20 healthy men were compared. Blood samples were collected at three time points: before treadmill running (pre-EX), immediately after treadmill running (post-EX), and at 60 min after treadmill running (60 min post-EX). Expression levels of irisin, FGF21, adiponectin, FFA, DA, and 5-HT were increased after 30 min of treadmill running exercise regardless of exercise intensity. Their levels were increased at 60 min post-EX. They showed no significant difference immediately after exercise regardless of exercise intensity. Only irisin, FGF21, FFA, and DA levels showed significant differences between moderate-intensity (50% VO_2_max) group and high-intensity group (80% VO_2_max) at 60 min post-EX. Considering that FFA level at 80% VO_2_max was significantly lower than that at 50% VO_2_max during post-EX and side effects of high-intensity exercise, moderate-intensity exercise would be a more competitive method compared to high-intensity for the prevention and improvement of obesity and obesity-related metabolic diseases.

## Introduction

Obesity and overweight as principal risk factors for mortality-relevant diseases by high body mass index (BMI) [1] have increased rapidly worldwide and amounted to pandemic ranges [2, 3]. They are linked to a variety of metabolic and chronic diseases, such as cardiovascular diseases, insulin resistance, type 2 diabetes mellitus (T2DM), respiratory diseases, fatty liver diseases, and certain types of cancers [2, 4]. Since overweight and obesity are preventable, continuous research and intervention are required for their prevention [2, 3].

Exercise, including physical activity is recognized as a means for maintaining health and for treating various life-threatening diseases such as obesity-induced diseases. Although further research is needed to elucidate the clear mechanism by which exercise has beneficial effects on the human body, this mechanism is undoubtedly mediated by alteration of neurotransmitters and peptides levels induced by physical activity. Such exercise-induced health-aiding results can protect from numerous obesity-induced diseases possible due to exercise-induced release of cytokines (including myokines, adipokines, and hepatokines) such as irisin, fibroblast growth factor-21 (FGF21), and adiponectin [5].

Irisin is an exercise-induced adipo-myokine. It plays a key role in converting white adipose tissue (WAT) into brown adipose tissue (BAT) through browning [6-8]. Irisin is also expected to play a potential role in improving obesity and obesity-related metabolic conditions [9, 10] because various studies have shown that irisin can improve glucose and lipid metabolism [11, 12], reduce weight and enhance glucose homeostasis [7, 13], improve insulin resistance [14, 15] and raises oxidative metabolism in cells [16].

Similarly, FGF21 is also an exercise-induced hepatokine. It is a member of the endocrine FGF family that plays and important role in metabolic homeostasis [17]. FGF21 also plays a role in converting WAT into brown adipocytes [18]. In addition, FGF21 and FGF21 analogues can improve insulin sensitivity, improve glucose and lipid metabolism, reduce hepatosteatosis, and increase energy expenditure [19-21]. Therefore, FGF21 has attracted attention as a treatment for obesity and metabolic diseases [22].

Increased level of adiponectin is one of mechanisms by which exercise can induce obesity-related metabolic improvements [23, 24]. The effect of adiponectin in improving obesity and obesity-related diseases has been well established [25, 26].

Dopamine (DA) and serotonin (5-HT) are related to fatigue of the central nervous system (CNS) during exercise [27], and obesity due to diet control [28, 29] It has been suggested that DA can associated control food intake [30-32], although this remains controversial. In addition, the appetite-regulating effect of 5-HT related to satiety in the process of food intake has long been demonstrated [33-35].

Many studies to date have reported changes in cytokines and neurotransmitter levels caused by exercise. Our previous study has confirmed that a single bout of acute exercise can significantly increase FGF21 levels in both rats and human with significant increases of free fatty acid (FFA) but significant decreases of insulin levels, suggesting that exercise is a therapeutic strategy for diabetes and obesity [36]. In addition, it has been confirmed that 75% of maximal oxygen uptake (VO_2_max) treadmill running for 40 min can increase DA and 5-HT levels regardless of caffeine intake [37]. However, levels of these neurotransmitter and cytokines can be affected by the intensity of exercise.

The intensity of exercise may an important variable to explain health benefits of physical activity. Change in levels of cytokines and neurotransmitters according to exercise intensity can associated with health benefits induced by exercise. In other words, investigating levels of neurotransmitters and cytokines according to exercise intensity could suggest a way to maximize health benefits of exercise. Therefore, the objective of this study was to the effectiveness of exercise by checking changes in expression levels of irisin, FGF21, DA, 5-HT, adiponectin, and FFA after 50% VO_2_max and 80% VO_2_max treadmill running for 30 min and present a relatively competitive protocol for preventing obesity and related metabolic diseases.

## Material and methods

### Subjects

Twenty healthy male non-athletic volunteers participated in this study (age, 23.64 ± 3.18 years; height, 174.4 ± 4.69 cm; weight, 69.72 ± 5.08 kg; body surface area (BSA), 1.82 ± 0.2 m^2^; BMI, 22.4 ± 2.85 kg/m^2^; Body fat, 20.62 ± 2.57%; VO_2_max, 42.36 ± 4.61 ml⋅kg^−1^⋅min^−1^). Physical characteristics of these study subjects are shown in Table 1. Body fat percentage was measured with the bio-impedance method (Inbody 770, Seoul, Korea). Body surface area was calculated according to the Du Bois formula [38].

**Table 1.** Physical characteristics of subjects.

| Variables (unit) | Mean ± SD |
| --- | --- |
| Age (yrs) | 23.64±3.18 |
| Height (cm) | 174.40±4.69 |
| Weight (kg) | 69.72±5.08 |
| BSA (m<sup>2</sup>) | 1.82±0.21 |
| BMI (kg/m<sup>2</sup>) | 22.92±2.85 |
| Body fat (%) | 23.62±2.57 |
| VO<sub>2</sub>max (ml·kg<sup>-1</sup>·min<sup>-1</sup>) | 42.36±4.61 |

BSA, body surface area; BMI, body mass index; VO<sub>2</sub>max, max oxygen uptake.

Inclusion criteria were as follows: 1) absence of health problems, and 2) no smoking history. Each subject provided written informed consent to participate in this study, after the purpose, experimental procedures, and any potential risks were explained. All subjects read and signed an informed consent regarding the purpose of this study and procedures. They were asked to refrain from alcohol consumption, smoking, medication, and vigorous physical activity during 48 hours before the testing. All experimental protocols were approved by Soonchunhyang University Research Committee. Procedures complied with the 2013 Declaration of Helsinki of the World Medical Association. Studies involving human participants were reviewed and approved by the Institutional Review Board on Human Subjects Research and Ethics Committees, Soonchunhyang University (No. 1040875-201611-BR-042 and No. 1040875-202104-BR-030).

### Measurement and experimental procedures

This study was conducted in a climate chamber between 2–5 p.m. to control for the influence of body temperature circadian rhythm. Environmental conditions were maintained at a temperature of 24.5 ± 0.3°C, a relative humidity of 50 ± 3.0%, and an air velocity of 1 m/sec.

Regarding exercise conditions and monitoring for human subjects, the subject sat in a chair in a relaxed posture for 30 min. Later, 50% VO_2_max treadmill running and 80% VO_2_max treadmill running (Quinton Medtrack SR 60) exercises were conducted for 30 min without drinking. After finishing the treadmill running exercise, a drinkable 500 mL of tepid water bottle was provided to each subject to prevent dehydration. One week prior to the treadmill test, physical load (VO_2_max) was determined by performing prolonged running on a treadmill (gradually increased from 2 to 16 km/h) until the subject became exhausted. Twenty subjects underwent a treadmill running for 30 min at 50% and 80% VO_2_max measured with an expired air gas analyzer (Quark Pulmonary Function Testing Lung Volumes Module 2 ergo, COSMED). All subjects performed 50% VO_2_max treadmill running first. They then performed a treadmill running at 80% VO_2_max five days later.

### Blood analysis

Blood was sampled before treadmill running exercise (pre-EX), immediately after 30 min of treadmill running exercise (post-EX), and at 60 min of rest after 30 min of treadmill running exercise (60 min post-EX). Schematic 1 shows the experiment protocol. Blood samples were collected from the antecubital vein and left at room temperature for 15−30 min to enable clotting, after which samples were centrifuged at 3,000 rpm (2,000 × g) for 10 min at 4°C. Serum was then harvested and stored at −80°C until analyzed. Irisin was determined using a commercial Enzyme-Linked Immunosorbent Assay (ELISA) kit (Irisin EIA kit EK-067-52;Phoenix Pharmaceuticals, Burlingame, CA, USA). FGF21 was measured using a Human FGF21 Quantikine ELISA Kit (R&D Systems). Adiponectin level was measured using an ELISA kit (Human Adiponectin ELISA, Zell BioGmbH, Ulm, Germany). FFA level was measured using an automated biochemical analyzer (Hitachi 7180). 5-HT level was determined via a high-pressure liquid chromatography (HPLC) using an Alliance Waters 465 HPLC (Waters, USA) in combination with a Serotonin Kit (Recipe, Munich, Germany). Blood samples were collected into EDTA tubes from antecubital veins. These samples were centrifuged at 3000 rpm (2000 x g) for 10 min at 4°C. Plasma was harvested and stored at −80°C until analyzed. DA level was determined using the HPLC method (Plasma Catecholamine Kit, Bio-Rad, Germany) in combination with an Acclaim HPLC (Bio-Rad, USA).

**Schematic 1.**
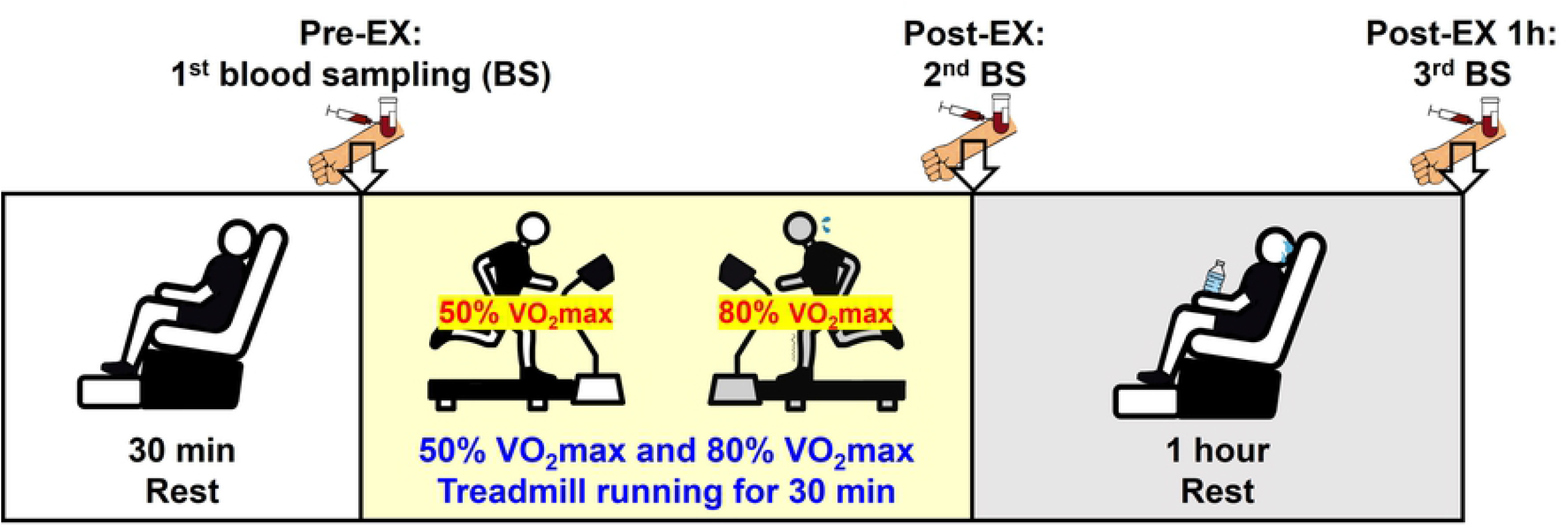
Schematic showing the protocol of experiments. Blood sampling (BS) was done before starting the treadmill running exercise (Pre-EX), immediately after treadmill running (50% VO_2_max and 80% VO_2_max) exercise for 30 min (Post-EX), and at 60 min of rest after treadmill running (50% VO_2_max and 80% VO_2_max) exercise for 30 min (60 min Post-EX).

### Statistical analysis

Descriptive statistics are expressed as mean ± standard deviation (SD) using a commercially available computer software SPSS for windows, version 21.0 (SPSS Inc., Chicago, IL, USA). Wilcoxon signed rank test and Mann−Whitney U-test were used to examine differences between groups and within groups. Significant differences were considered at *p <* 0.05.

## Results

### Irisin level

Post-EX irisin levels were significantly higher than pre-EX irisin levels after both moderate-intensity exercise and high-intensity exercise (50% VO_2_max treadmill running, 7.32 ± 3.09 vs. 8.74 ± 3.85 ng/mL, *p <* 0.01; 80% VO_2_max treadmill running, 7.28 ± 3.06 vs. 9.35 ± 3.92 ng/mL, *p <* 0.01). However, irisin levels were not significantly different between the two exercise groups (50% VO_2_max post-EX and 80% VO_2_max post-EX) (Figure 1). Moreover, irisin levels at 60 min post-EX were significantly higher than post-EX levels of irisin after exercise regardless of exercise intensity (50% VO_2_max treadmill running, 8.74 ± 3.85 vs. 9.86 ± 4.07 ng/mL, *p <* 0.001; 80% VO_2_max treadmill running, 9.35 ± 3.92 vs. 11.24 ± 4.51 ng/mL, *p <* 0.001). At 60 min post-EX, irisin levels in the high-intensity exercise (80% VO_2_max) group were significantly higher than those in the moderate-intensity exercise (50% VO_2_max) group (*p <* 0.05).

**Figure 1.**
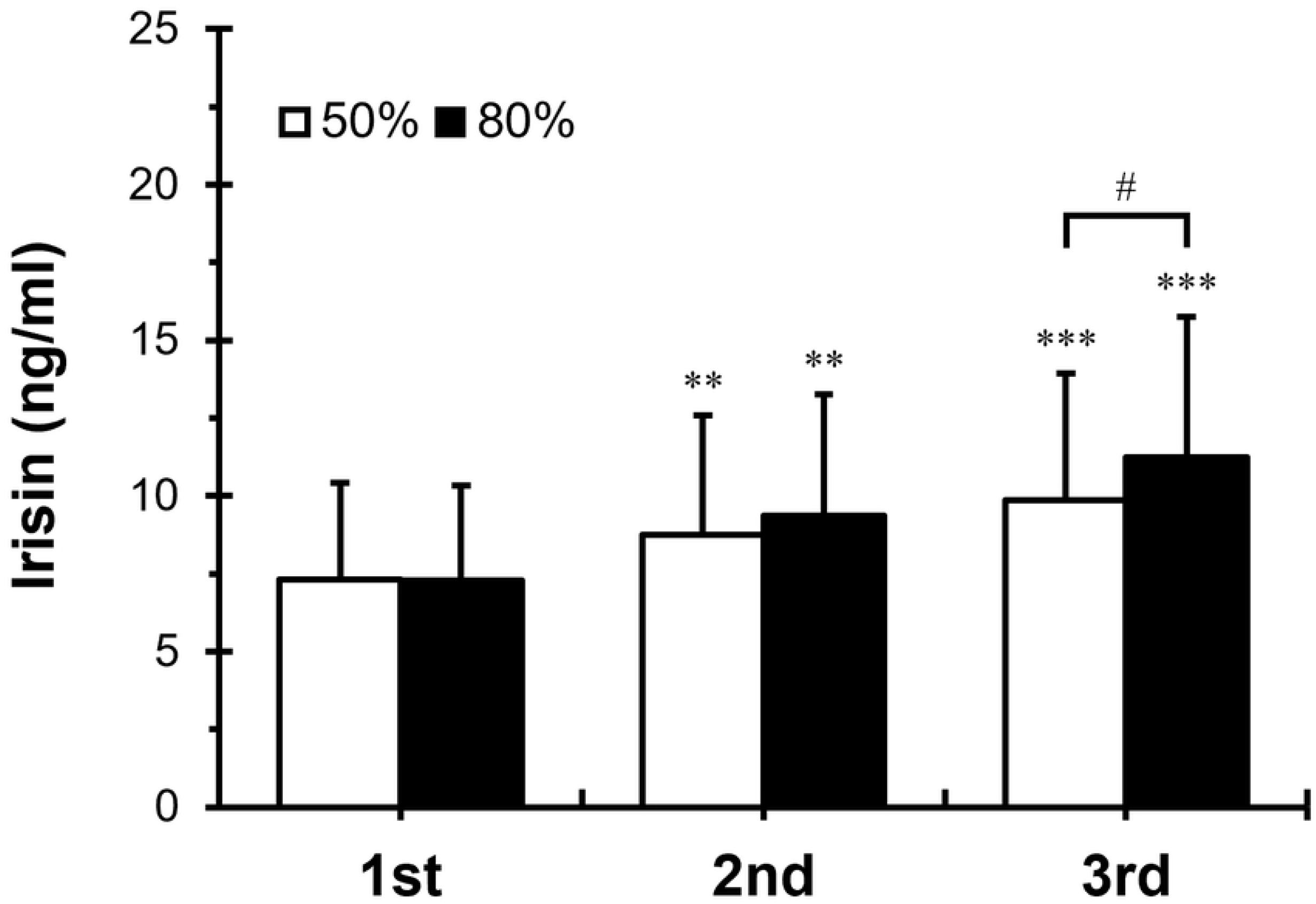
Blood irisin concentrations at Pre-exercise, Post-exercise, and 60 min post-exercise. 1st, before treadmill running exercise; 2nd, immediately after treadmill running exercise at 50% and 80% VO_2_max for 30 min; 3rd, 60 min after treadmill running exercise at 50% and 80% VO_2_max for 30 min. White columns indicate treadmill running exercise at 50% VO_2_max for 30 min. Black columns indicate treadmill running exercise at 80% VO_2_max for 30 min. Values are presented as mean ± SD. Significant difference, 50% versus 80% VO_2_max, ^*#*^*P <* 0.05. Significant difference, 1st versus 2nd; 2nd versus 3rd, *\*\*, P <* 0.01; and *\*\*\*, P <* 0.001.

### FGF21 level

FGF21 levels post-EX were significantly higher than pre-EX FGF21 levels after exercise regardless of exercise intensity (50% VO_2_max treadmill running, 37.54 ± 22.81 vs. 47.03 ± 24.28 pg/mL, *p <* 0.001; 80% VO_2_max treadmill running, 38.16 ± 20.75 vs. 49.55 ± 26.91 pg/mL, *p <* 0.001). However, FGF21 levels post-EX were not significantly different between 50% VO_2_max and 80% VO_2_max exercise groups (Figure 2). Moreover, FGF21 levels at 60 min post-EX were significantly higher than their post-EX levels in both exercise groups (50% VO_2_max treadmill running, 47.03 ± 24.28 vs. 64.20 ± 31.27 pg/mL, *p <* 0.001; 80% VO_2_max treadmill running, 49.55 ± 26.91 vs. 72.36 ± 28.42 pg/mL, *p <* 0.001). At 60 min post-EX, FGF21 levels in the high-intensity exercise (80% VO_2_max) group were significantly higher than those in the moderate-intensity (50% VO_2_max) group (*p <* 0.05).

**Figure 2.**
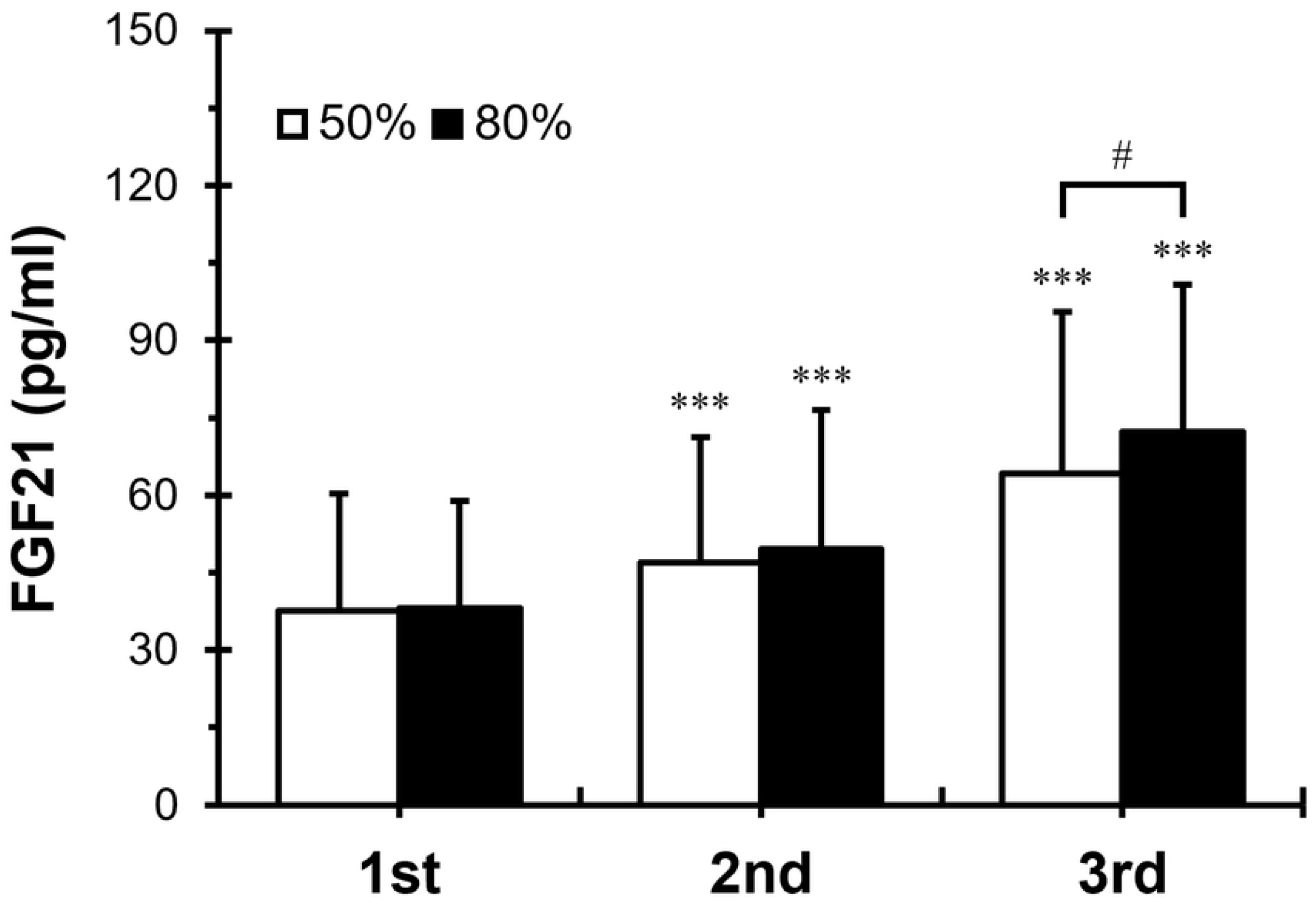
Blood FGF21 concentrations at Pre-exercise, Post-exercise, and 60 min post-exercise. 1st, before treadmill running exercise; 2nd, immediately after treadmill running exercise at 50% and 80% VO_2_max for 30 min; 3rd, 60 min after treadmill running exercise with 50% and 80% VO_2_max for 30 min. White columns indicate treadmill running exercise at 50% VO_2_max for 30 min. Black columns indicate treadmill running exercise at 80% VO_2_max for 30 min. Values are presented as mean ± SD. Significant difference, 50% versus 80% VO_2_max, ^*#*^ *P <* 0.05. Significant difference of the 1st versus the 2nd; 2nd versus 3rd, *\*\*\*, P <* 0.001. FGF21, fibroblast growth factor-21.

### Adiponectin level

Post-EX adiponectin levels were significantly higher than pre-EX adiponectin levels in both exercise groups (50% VO_2_max treadmill running, 8.02 ± 4.36 vs. 8.87 ± 5.21 μg/mL, *p <* 0.01; 80% VO_2_max treadmill running, 8.09 ± 4.29 vs. 9.03 ± 4.86 μg/mL, *p <* 0.01). However, post-EX adiponectin levels were not significantly different between the two exercise groups (50% VO_2_max post-EX and 80% VO_2_max post-EX) (Figure 3). Moreover, adiponectin levels at 60 min post-EX were significantly higher than their post-EX levels in both exercise groups (50% VO_2_max treadmill running, 8.87 ± 5.21 vs. 10.78 ± 5.63 μg/mL, *p <* 0.001; 80% VO_2_max treadmill running, 9.03 ± 4.86 vs. 11.23 ± 5.58 μg/mL, *p <* 0.001). However, adiponectin levels at 60 min post-EX were not significantly different between the two exercise groups (50% VO_2_max 60 min post-EX and 80% VO_2_max 60 min post-EX).

**Figure 3.**
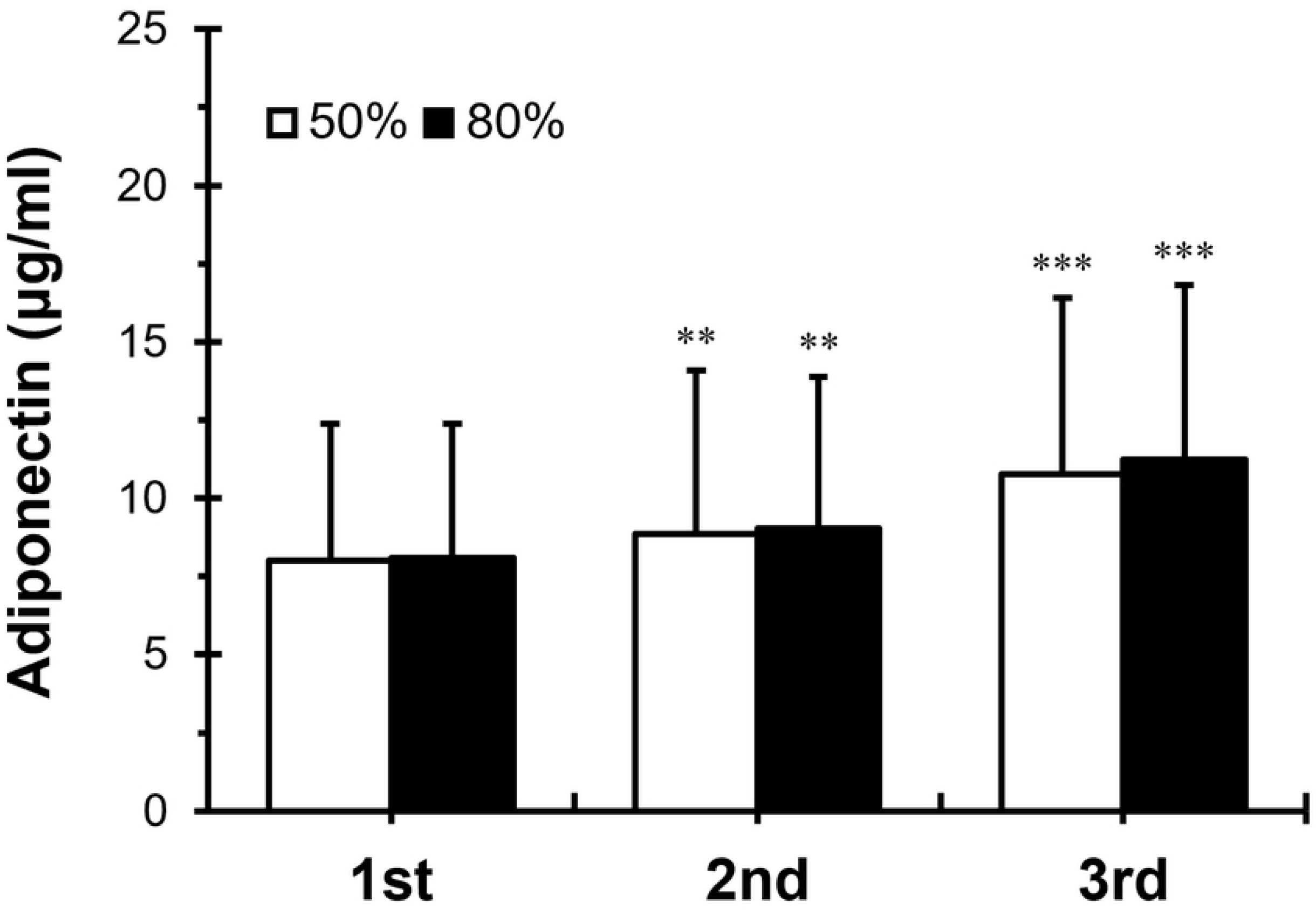
Blood adiponectin concentrations at Pre-exercise, Post-exercise, and 60 min post-exercise. 1st, before treadmill running exercise; 2nd, immediately after treadmill running exercise at 50% and 80% VO_2_max for 30 min; 3rd, 60 min after treadmill running exercise at 50% and 80% VO_2_max for 30 min. White columns indicate treadmill running exercise at 50% VO_2_max for 30 min. Black columns indicate treadmill running exercise at 80% VO_2_max for 30 min. Values are presented as means ± SD. Significant difference, 1st versus 2nd; 2nd versus 3rd, *\*\*, P <* 0.01 and *\*\*\*, P <* 0.001.

### FFA level

Post-EX FFA levels were significantly increased compared with pre-EX FFA levels in both exercise groups (50% VO_2_max treadmill running, 457.28 ± 97.72 vs. 534.25 ± 102.35 nEq/mL, *p <* 0.05; 80% VO_2_max treadmill running, 451.97 ± 95.62 vs. 512.19 ± 106.28 nEq/mL, *p <* 0.05.) However, post-EX FFA levels were not significantly different between the two exercise groups (50% VO_2_max post-EX and 80% VO_2_max post-EX) (Figure 4). Moreover, FFA levels at 60 min post-EX were significantly higher than post-EX FFA levels in both exercise groups (50% VO_2_max treadmill running, 534.25 ± 102.35 vs. 584.57 ± 129.05 nEq/mL, *p <* 0.001; 80% VO_2_max treadmill running, 512.19 ± 106.28 vs. 532.61 ± 127.38 nEq/mL, *p <* 0.001). At 60 min post-EX, FFA levels in the high-intensity exercise (80% VO_2_max) group were significantly lower than those in the moderate-intensity exercise (50% VO_2_max) group (*p <* 0.05).

**Figure 4.**
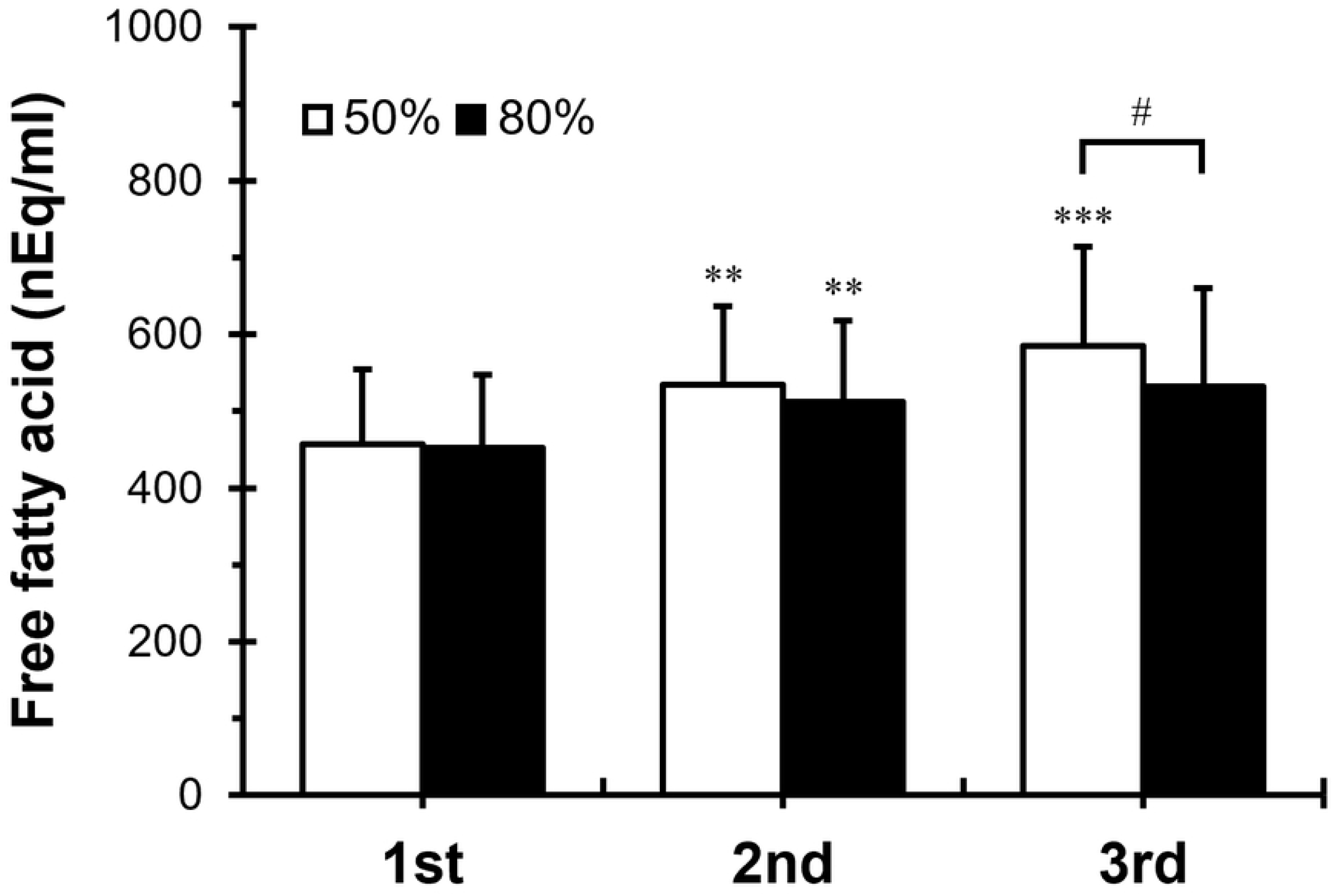
Blood Free fatty acid concentrations at Pre-exercise, Post-exercise, and 60 min post-exercise. 1st, before treadmill running exercise; 2nd, immediately after treadmill running exercise at 50% and 80% VO_2_max for 30 min; 3rd, 60 min after treadmill running exercise at 50% and 80% VO_2_max for 30 min. White columns indicate treadmill running exercise at 50% VO_2_max for 30 min. Black columns indicate treadmill running exercise at 80% VO_2_max for 30 min. Values are presented as mean ± SD. Significant difference, 50% versus 80% VO_2_max, ^*#*^*P <* 0.05. Significant difference, 1st versus 2nd; 2nd versus 3rd, *\*\*, P <* 0.01 and *\*\*\*, P <* 0.001.

### Dopamine level

Post-EX DA levels were significantly higher than pre-EX DA levels in both exercise groups (50% VO_2_max treadmill running, 9.72 ± 5.07 vs. 12.63 ± 7.10 pg/mL, *p <* 0.001; 80% VO_2_max treadmill running, 9.59 ± 4.98 vs. 13.26 ± 8.07 pg/mL, *p <* 0.001). However, post-EX DA levels were not significantly different between the two exercise groups (50% VO_2_max post-EX and 80% VO_2_max post-EX) (Figure 5). Moreover, DA levels at 60 min post-EX were significantly higher than post-EX DA levels in both exercise groups (50% VO_2_max treadmill running, 12.63 ± 7.10 vs. 19.22 ± 9.30 pg/mL, *p <* 0.001; 80% VO_2_max treadmill running, 13.26 ± 8.07 vs. 23.28 ± 9.17 pg/mL, *p <* 0.001). At 60 min post-EX, DA levels in the high-intensity exercise (80% VO_2_max) group were significantly higher than those in the moderate-intensity exercise (50% VO_2_max) group (*p <* 0.05).

**Figure 5.**
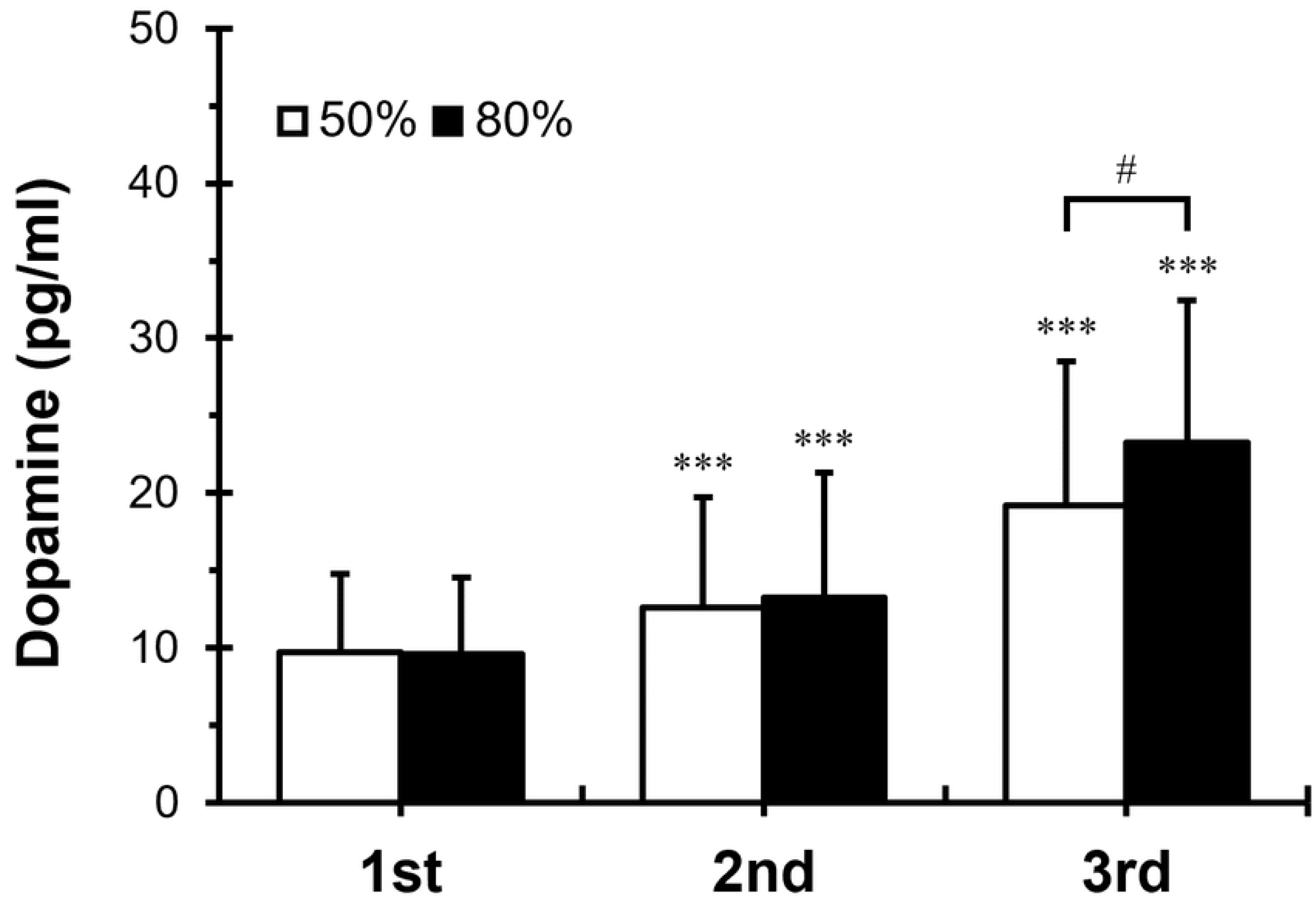
Blood dopamine concentrations at Pre-exercise, Post-exercise, and 60 min post-exercise. 1st, before treadmill running exercise; 2nd, immediately after treadmill running exercise at 50% and 80% VO_2_max for 30 min; 3rd, 60 min after treadmill running exercise at 50% and 80% VO_2_max for 30 min. White columns indicate treadmill running exercise at 50% VO_2_max for 30 min. Black columns indicate treadmill running exercise at 80% VO_2_max for 30 min. Values are presented as mean ± SD. Significant difference, 50% versus 80% VO_2_max, ^*#*^*P <* 0.05. Significant difference, 1st versus 2nd; 2nd versus 3rd, *\*\*\*, P <* 0.001.

**Figure 6.**
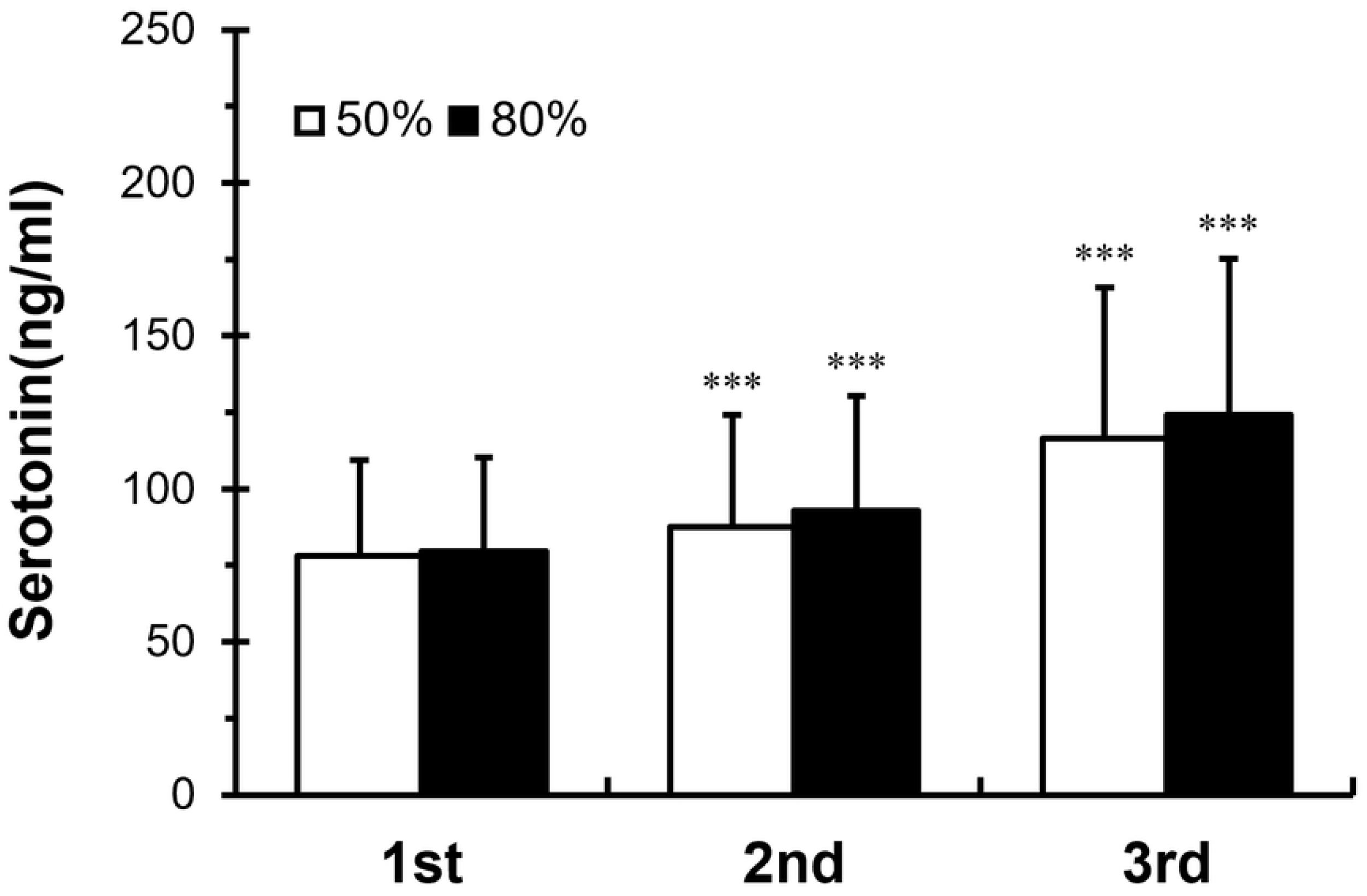
Blood serotonin concentrations at Pre-exercise, Post-exercise, and 60 min after post-exercise. 1st, before treadmill running exercise; 2nd, immediately after treadmill running exercise at 50% and 80% VO_2_max for 30 min; 3rd, 60 min after treadmill running exercise at 50% and 80% VO_2_max for 30 min. White columns indicate treadmill running exercise at 50% VO_2_max for 30 min. Black columns indicate treadmill running exercise at 80% VO_2_max for 30 min. Values are presented as mean ± SD. Significant difference, 1st versus 2nd; 2nd versus 3rd, *\*\*\*, P <* 0.001.

### Serotonin level

Post-EX 5-HT levels were significantly higher than pre-EX 5-HT levels after exercise regardless of intensity (50% VO_2_max treadmill running, 78.16 ± 31.06 vs. 84.55 ± 36.45 ng/mL, *p <* 0.001; 80% VO_2_max treadmill running, 79.41 ± 30.82 vs. 85.91 ± 37.54 ng/mL, *p <* 0.001). However, post-EX 5-HT levels were not significantly different between the two exercise groups (50% VO_2_max treadmill running, 84.55 ± 36.45 vs. 116.47 ± 49.38 ng/mL, *p <* 0.001; 80% VO_2_max treadmill running, 85.91 ± 37.54 vs. 124.05 ± 51.25 ng/mL, *p <* 0.001). However, at 60 min post-EX, 5-HT levels were not significantly different between the two exercise groups (50% VO_2_max 60 min post-EX and 80 VO_2_max 60 min post-EX).

## Discussion

This study investigated effects of two different intensity exercises [treadmill running exercise for 30 min at moderate intensity (50% VO_2_max) and high-intensity (80% VO_2_max)] on expression levels of irisin, FGF21, adiponectin, FFA, DA, and 5-HT. More specifically, to suggest a relatively competitive exercise protocol for preventing and improving obesity and obesity-related metabolic diseases, levels of irisin, FGF21, adiponectin, FFA, DA, and 5-HT after moderate-intensity (50% VO_2_max) and high-intensity (80% VO_2_max) treadmill running exercise were examined at three time points: before (pre-EX), immediately after (post-EX), and at 60 min after the exercise (60 min post-EX). Only statistically significant differences were found for levels of irisin, FGF21, FFA, and DA at 60 min post-EX between the two exercise groups.

Physical activity can promote health and lower the incidence of multifarious diseases over numerous molecular pathways and cytokines such as myokines, adipokines, and hepatokines [39]. Irisin can increase browning of WAT [6-8], increase glucose uptake [11], improve fatty acid oxidation [12], increase lipolysis [40], improve insulin resistance [14, 15], decrease body weight [7, 13], and improve metabolism [9, 40]. Thus, it is expected to be helpful in improving and preventing obesity and obesity-related metabolic diseases [9, 10] [Schematic 2]. As such, modulating circulation irisin levels can aid in the prevention of various endocrine and metabolic diseases [41]. Irisin expression affected by physical activity has been previously debatable. However, it has recently recognized that irisin can be induced by exercise. Results of our study showed that irisin levels post-EX and at 60 min post-EX were significantly increased than pre-EX irisin levels regardless of exercise intensity, consistent with studies reporting an increase of irisin level after exercise [11, 42, 43]. Different from previous results showing that irisin levels the moderate-intensity exercise can be higher than high-intensity exercise [42], our study showed higher expression of irisin during high-intensity exercise, which was higher at 60 min post-EX was only significant. Our results showed higher irisin levels at 60 min post-EX than post-EX irisin levels, not consistent with one study showing recovering to resting levels of irisin at 60 min after exercise [44]. It may suggest that, in contrast with our study of 30 min of treadmill running at 50% VO_2_max and 80% VO_2_max, irisin levels might have sharply increased after a relatively high-intensity exercise such as one that leads to just exhaustion, resulting in a faster recovery to baseline levels.

**Schematic 2.**
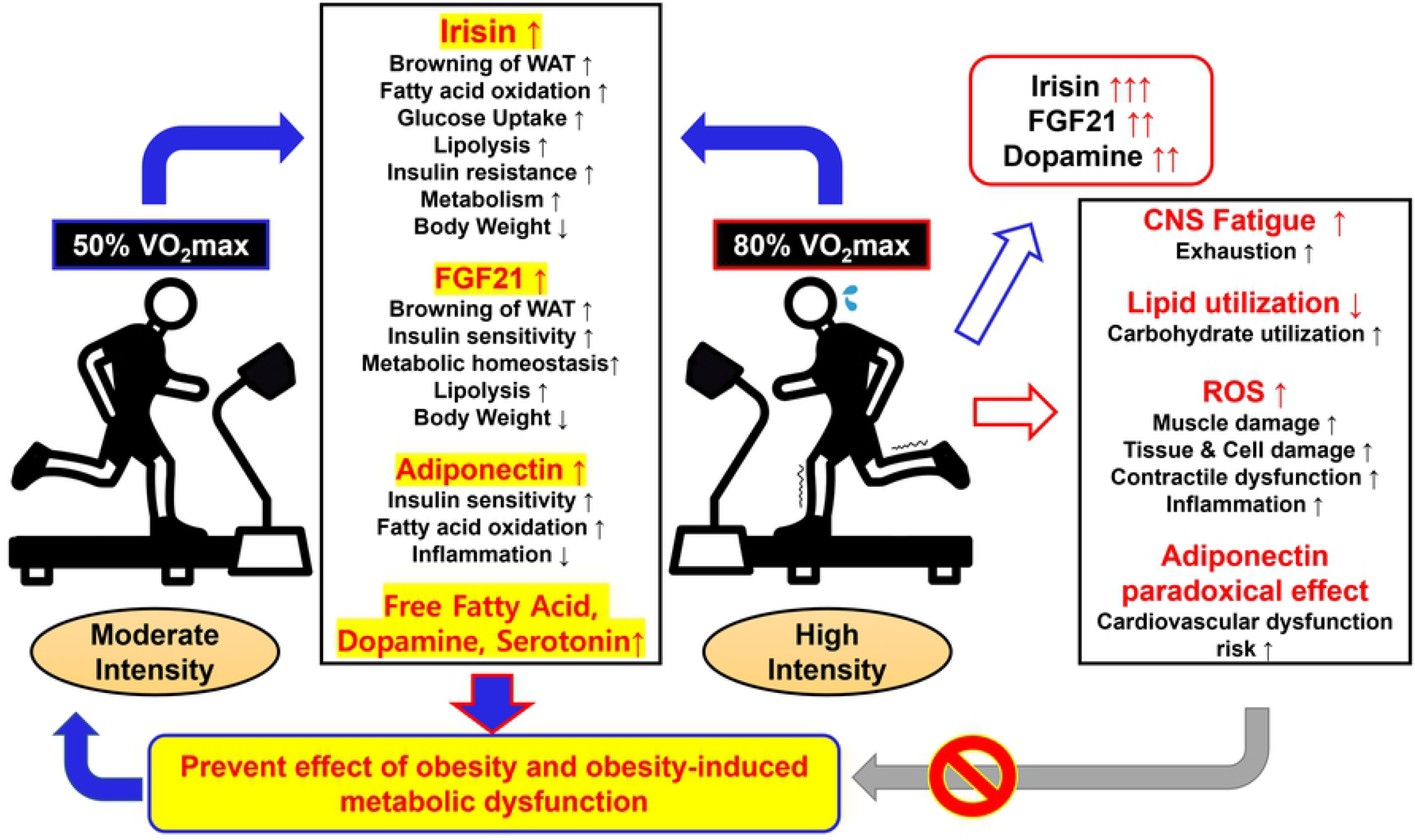
Summary of effects of 50% and 80% VO_2_max treadmill running exercise on obesity and obesity-induced metabolic dysfunction exercise. ↑ indicates increase or improvement; ↓ indicates decrease. WAT, white adipose tissue; FGF21, fibroblast growth factor-21; CNS, central nervous system; ROS, reactive oxygen species.

FGF21 may maintain energy homeostasis and improve metabolic diseases and obesity [45]. FGF21 can increase browning of WAT [18, 46-48], improve insulin sensitivity [19, 21, 48], increase energy expenditure [19, 46, 48], improve metabolic homeostasis [20, 49, 50], decrease body weight [19, 48, 49], and inhibit lipid accumulation [51]. It may be useful for the improvement and prevention of obesity and obesity-related metabolic diseases [Schematic 2]. Exercise intensity may decide FGF21 expression levels and recover side effects of obesity in subcutaneous WAT [52]. Consistent with the present study, several studies have reported significant increases of FGF21 levels after acute exercise [53, 54] However, there might be differences in the timing at which the expression of FGF21 is induced by exercise. In our previous study, FGF21 level between not significantly pre-EX and post-EX at 50% and 80% VO_2_max while at 60 min post-EX, FGF21 level was significantly higher than pre-EX [36]. In a recent study, FGF21 level was significantly higher after 60 min from 55% and 75% VO_2_peak treadmill running than before treadmill running but between before and immediately after significantly at 55% VO_2_peak treadmill running however not 75% VO_2_peak [53]. This difference in FGF21 of expression might be due to different characteristics of participants or differences in study designs since FGF21 levels induced by the same exercise might be affected by characteristics of participants, such as weight, BMI, age, and disease status [54, 55]. Nevertheless, most reports have reported that acute exercise can induce the expression of FGF21, consistent with our studies. The level of FGF21 has been consistently shown to be higher in a high-intensity group than that in a moderate-intensity group, especially at 60 min after the exercise [36, 53]. Interestingly, at 60 min after exercise, the high-intensity exercise group showed significantly higher levels of FGF21 than the moderate-intensity exercise group, although significant differences in physiological responses such as plasma glucose and insulin levels were not found between the two groups [53].

Adiponectin can improve insulin sensitivity and insulin resistance [56, 57], increase fatty acid oxidation [57], and decrease inflammation [58, 59]. Thus, it can exert positive effects on obesity and obesity-related metabolic diseases [Schematic 2].

Although the effect of exercise on adiponectin levels may vary depending on exercise intensity, duration, acute or chronic, and body fat mass, exercise is associated with an increase of adiponectin level [60, 61]. Consistently, our study found that 30 min of treadmill running exercise could significantly increase adiponectin levels regardless of exercise intensity. However, data for demonstrating that acute exercise can increase adiponectin levels are insufficient [62]. Thus, further studies are needed.

The secretion and synthesis of neurotransmitters such as 5-HT and DA are affected by exercise intensity [63]. We found that treadmill running exercise increased DA and 5-HT levels, consistent with our previous studies [37]. DA levels were significantly higher in the high-intensity exercise (80% VO_2_max) group than in the moderate-intensity exercise (50% VO_2_max) group. 5-HT levels also showed a trend to be higher in the high-intensity exercise (80% VO_2_max) group than in the moderate-intensity exercise (50% VO_2_max) group. However, the difference between the two groups was not significant. DA is involved in circuits related to reward and motivation. DA levels may weaken or strengthen control circuits of food intake regulation. In a recent study, exercise increased DA levels in the nucleus accumbens to aid facultative restoration of the reward system and decreased high-fat diet preference and energy efficiency in obese rats, demonstrating improvements of weight increase control [64].

5-HT has long been proven to have an appetite-regulating effect related to satiety during food intake and recognized as a treatment for obesity [34]. In addition, it should be noted that, since an increase in 5-HT levels during exercise is associated with the development of central fatigue, a higher intensity and prolonged exercise may lead to rapid fatigue and exhaustion due to a higher increase of 5-HT levels [27].

As a result of our study, levels of irisin, FGF21, and DA known to be effective in improving and preventing obesity and obesity-related metabolic diseases were more expressed after a high-intensity exercise than those after a moderate-intensity exercise. However, adiponectin and 5-HT levels were not significantly affected by exercise intensity.

FFA levels were found to be increased significantly at 60 min post-EX after a moderate-intensity exercise rather than after a high-intensity exercise. High-intensity exercise does not show more peripheral lipolysis than a moderate-intensity exercise [65]. Fat oxidation climaxes at moderate-intensity exercise (45-65% VO_2_max), and more high-intensity exercise lowers fat oxidation [66]. During low- and moderate-intensity exercise, lipids are mainly used as energy but, during a high-intensity exercise, the use of lipids is sharply reduced while the utilization of carbohydrates is increased [67] [Schematic 2]. Consistently, our study results revealed that post-EX FFA levels tended to be higher (although not statistically significant) after a moderate-intensity exercise than those after a high-intensity exercise and that FFA levels were significantly higher at 60 min post-EX after a moderate-intensity exercise.

Nevertheless, other authors reviewed that high-intensity exercise can lead to pathological conditions in multiple organs beyond performance degradation due to high-intensity exercise-induced other cytokines expression [68]. Thus, when presenting exercise intensity for improvement and prevention of obesity and obesity-related metabolic diseases, side effects of high-intensity exercise should be considered.

Higher intensity exercise can result in significantly higher reactive oxygen species (ROS), which induce tissue damage [69, 70]. High levels of ROS from a high-intensity exercise may have harmful effects on cells, increase the risk of aging and age-related diseases, lead to contractile dysfunction, and result in muscle inflammation and damage [69-71]. In addition, higher levels of adiponectin resulting from a high-intensity exercise may play a detrimental role in cardiovascular function, similar to lower levels of adiponectin [72] [Schematic 2].

Since these side effects of a high-intensity exercise may increase the risk of health conditions, it is considered that it is inappropriate to perform 80%VO2max exercise for high expression of irisin, FGF21, and DA. Considering these side effects of a high-intensity exercise and that both 50%VO2max and 80%VO2max groups showed significant increases of irisin, FGF21, adiponectin, FFA, DA, and 5-HT levels for preventing obesity and obesity-related metabolic diseases, 50%VO2max exercise could be suggested as a more competitive method [Schematic 2].

However, obesity and obesity-related metabolic disorders-related factors (such as orexin, leptin, HbA1c, and insulin) after moderate and high-intensity exercise should be investigated in future studies. In addition, our study did not investigate factors related to side effects that might occur from high-intensity exercise. Thus, inflammation-related markers, lipid utilization, and muscle damage factors according to exercise intensity should be investigated in the future.

## ABBREVIATIONS

FGF21: fibroblast growth factor-21
FFA: free fatty acids
DA: dopamine
5-HT: serotonin
VO_2_max: max oxygen uptake

## Acknowledgments

The authors thank study subjects whose participation made this study possible. The authors have no conflicts of interest, financial or otherwise, to disclose. This research was supported by a grant (No.2016R1D1A3B02015394) of the Basic Science Research Program through the National Research Foundation (NRF) funded by the Ministry of Education, Republic of Korea.

